# CTCF is crucial for decidualization of uterine stromal cells through facilitating HOXA11 expression

**DOI:** 10.1101/2025.06.10.658981

**Authors:** Xiao-Qi Yang, Run-Fu Jiang, An-Nan Zhang, Yu-Ming Deng, Shi Tang, Qing-Yan Zhang, Yan-Wen Xu, Shi-Hua Yang, Ji-Long Liu

## Abstract

CCCTC-binding factor (CTCF), a highly versatile transcriptional regulator and chromatin architectural protein, plays a pivotal role in mammalian development. In this study, we investigated the function of CTCF in uterine biology during pregnancy. We observed that CTCF expression in the human endometrium decreases from the proliferative to the secretory phase of the menstrual cycle. Similarly, in cultured human endometrial stromal cells (hESCs), CTCF expression declines during in vitro decidualization. Using siRNA-mediated knockdown, we demonstrated that CTCF is essential for the early phase of decidualization but dispensable in later stages. Notably, CTCF depletion reduces the expression of HOXA11, a well-established regulator of decidualization, and overexpression of HOXA11 rescues the decidualization defects caused by CTCF loss, identifying HOXA11 as a key downstream effector of CTCF. Mechanistically, CTCF binds to the HOXA11 promoter to facilitate its transcriptional activation. Furthermore, uterine-specific knockout of CTCF in *Pgr*-Cre mice leads to female infertility, characterized by diminished HOXA11 expression, impaired uterine receptivity, and defective decidualization. Collectively, our findings highlight the critical role of the CTCF-HOXA11 axis in decidualization in both humans and mice, providing novel insights into the molecular regulation of this process and potential therapeutic targets for reproductive disorders.

## Introduction

The endometrium, the inner mucosal layer of the uterus, plays a pivotal role in human conception. This dynamic tissue undergoes cyclical regeneration, proliferation, differentiation, and desquamation hundreds of times during reproductive life, orchestrated by ovarian steroid hormones [1]. Structurally composed of fibroblast-like stromal cells and glandular epithelial cells, the endometrium executes decidualization—a specialized transformation program converting stromal fibroblasts into secretory decidual cells—that establishes the essential cellular environment for embryo implantation and subsequent placental development [2]. This tightly regulated process depends on postovulatory progesterone surges and localized cAMP-mediated signaling pathways [3–5]. Accumulating evidence implicates defective decidualization in the pathogenesis of recurrent pregnancy loss, preeclampsia, and endometriosis [1]. Consequently, unraveling the molecular mechanisms governing decidualization represents a critical translational imperative, with potential to inform therapeutic strategies for these prevalent reproductive disorders.

In mammalian genomes, the chromatin landscape is organized into sub-megabase-scale topologically associating domains (TADs) - three-dimensional structures characterized by preferential intra-domain interactions and sharply attenuated inter-domain contacts [6]. TAD boundaries are evolutionarily conserved and frequently marked by CCCTC-binding factor (CTCF) occupancy [7–9], a highly conserved 11-zinc-finger (ZF) DNA-binding protein [10]. Genomic profiling reveals that CTCF predominantly interacts with non-palindromic consensus motifs by ZFs 4-8 and 7-11 [11, 12]. Context-dependent functions enable CTCF to operate as a transcriptional regulator, activator, or chromatin insulator [13]. Constitutive CTCF deletion causes early embryonic lethality [14–16], while somatic conditional knockouts have uncovered critical roles in gametogenesis (spermatogenesis [17, 18] and oogenesis [19]) and embryonic organogenesis [20–22]. Recent work highlights its requirement for pubertal uterine maturation through regulation of progesterone-responsive genes (Ihh, Fst, Errfi1) [23]. However, the functional involvement of CTCF in decidualization and pregnancy maintenance remains uncharacterized.

Here, utilizing cultured human endometrial stromal cells (hESCs), we established the necessity of CTCF for in vitro decidualization. Murine models further corroborated its essential role in maintaining uterine functionality during implantation and decidualization processes. Mechanistic investigations revealed that CTCF occupancy at the HOXA11 promoter region is crucial for its transcriptional activation. Collectively, our findings substantiate CTCF as a critical regulator of decidualization through HOXA11 expression modulation.

## Results

### CTCF expression declines during decidualization

To explore CTCF expression dynamics in the human endometrium, we initially examined a publicly accessible microarray dataset (GSE4888) derived from endometrial biopsies [24]. This analysis uncovered a notable decline in CTCF mRNA levels from the proliferative to secretory phase of the menstrual cycle (**Fig. S1**). Immunohistochemical assessment of endometrial biopsies from healthy volunteers with regular cycles corroborated this temporal expression pattern, showing abundant CTCF protein expression in epithelial and stromal compartments during the proliferative phase, followed by marked downregulation in the secretory phase (**Fig. 1A**). To elucidate CTCF’s functional role during decidualization, we employed an in vitro model using primary human endometrial stromal cells (hESCs) treated with progesterone analog (MPA) and cAMP analog (8Br-cAMP). Decidualization was confirmed by the robust induction of canonical markers PRL and IGFBP1 (**Fig. 1B-C**). This differentiation process was accompanied by the suppression of CTCF expression at both transcriptional and translational levels (**Fig. 1B-D**).

**Figure 1.**
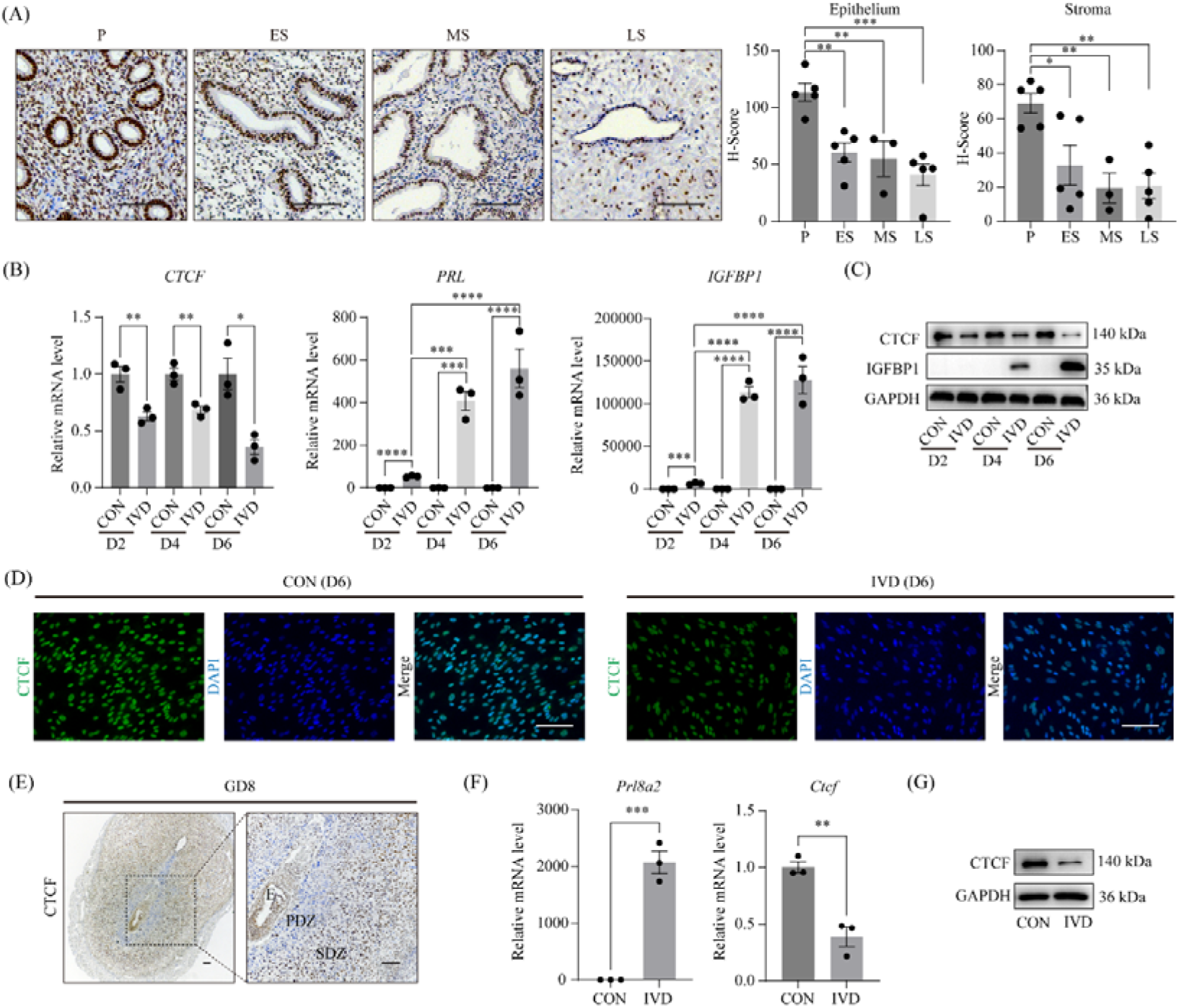
CTCF expression declines during decidualization. (A) Immunohistochemical analysis depicts CTCF protein expression in the endometrium across the menstrual cycle. P, proliferative phase; ES, early secretory phase; MS, middle secretory phase; LS, late secretory phase. Bar=100 μm. (B) Quantitative RT-PCR assesses *CTCF* RNA levels in hESCs during in vitro decidualizing at different time points. CON, vehicle control; IVD, in vitro decidualization. Data are presented as mean ± SEM. *, P < 0.05; **, P < 0.01; ***, P < 0.001; ****, P < 0.0001. (C-D) Western blot (C) and immunofluorescent staining (D) evaluate CTCF protein levels in hESCs during decidualization. Bar=100 μm. (E) Immunohistochemical staining shows CTCF protein in mouse uterus at the implantation site on gestational day 8 (GD8). E, embryo; PDZ, primary decidual zone; SDZ, secondary decidual zone. Bar=100 μm. (F-G) CTCF mRNA (F) and protein (G) expression in primary mouse endometrial stromal cells (mESCs) after in vitro decidualization. CON, vehicle control; IVD, in vitro decidualization. Data are presented as mean ± SEM. **, P < 0.01; ***, P < 0.001.

Notably, CTCF downregulation was also evident in the mouse uterus during pregnancy. Immunohistochemical analysis showed robust CTCF expression in stromal cells during the implantation window (GD4-5) (**Fig. S2A**), yet significantly reduced protein levels were noted in the primary decidual zone (PDZ) compared to the secondary decidual zone (SDZ) at GD8 (**Fig. 1E**). Spatiotemporal transcriptomic profiling further underscored progressive CTCF suppression in mesometrial decidual cells and decidua basalis throughout gestation (**Fig. S2B**). To corroborate these findings, we isolated primary mouse endometrial stromal cells (mESCs) from GD4 wild-type mice and induced decidualization in vitro. Consistent with hESC data, CTCF expression was markedly decreased during in vitro decidualization of mESCs (**Fig. 1F-G**). To elucidate the factors modulating CTCF expression, we individually treated hESCs with MPA or 8Br-cAMP and assessed their impact. Notably, MPA, but not 8Br-cAMP, significantly reduced CTCF expression (**Fig. S3A-D**). Collectively, these results suggest that P_4_ signaling mediates CTCF downregulation during decidualization.

### CTCF is indispensable for the early phase of decidualization but becomes dispensable in later stages

To evaluate CTCF’s role in hESCs, we conducted siRNA-mediated knockdown, validated by quantitative RT-PCR, western blot, and immunofluorescence (**Fig. S4A-C**). CTCF depletion impaired cell proliferation and migration without altering apoptosis (**Fig. S4D-G**). In vitro decidualization assays revealed significantly reduced PRL and IGFBP1 expression in CTCF-silenced hESCs following 4-day MPA plus 8-Br-cAMP treatment (**Fig. 2A-B**). Morphological analysis confirmed compromised decidualization, marked by diminished transition from fibroblastoid to epithelioid morphology (**Fig. 2C**). Temporal knockdown experiments demonstrated that CTCF suppression at day 2 of decidualization decreased IGFBP1 levels, whereas day 4 knockdown unexpectedly increased IGFBP1 expression (**Fig. 2D-E**). Conversely, CTCF overexpression prior to decidualization induction suppressed PRL and IGFBP1 expression (**Fig. 2F-G**). These results underscore CTCF’s pivotal role in initiating decidualization and the necessity for precise expression regulation, as both deficiency and excess disrupt proper differentiation.

**Figure 2.**
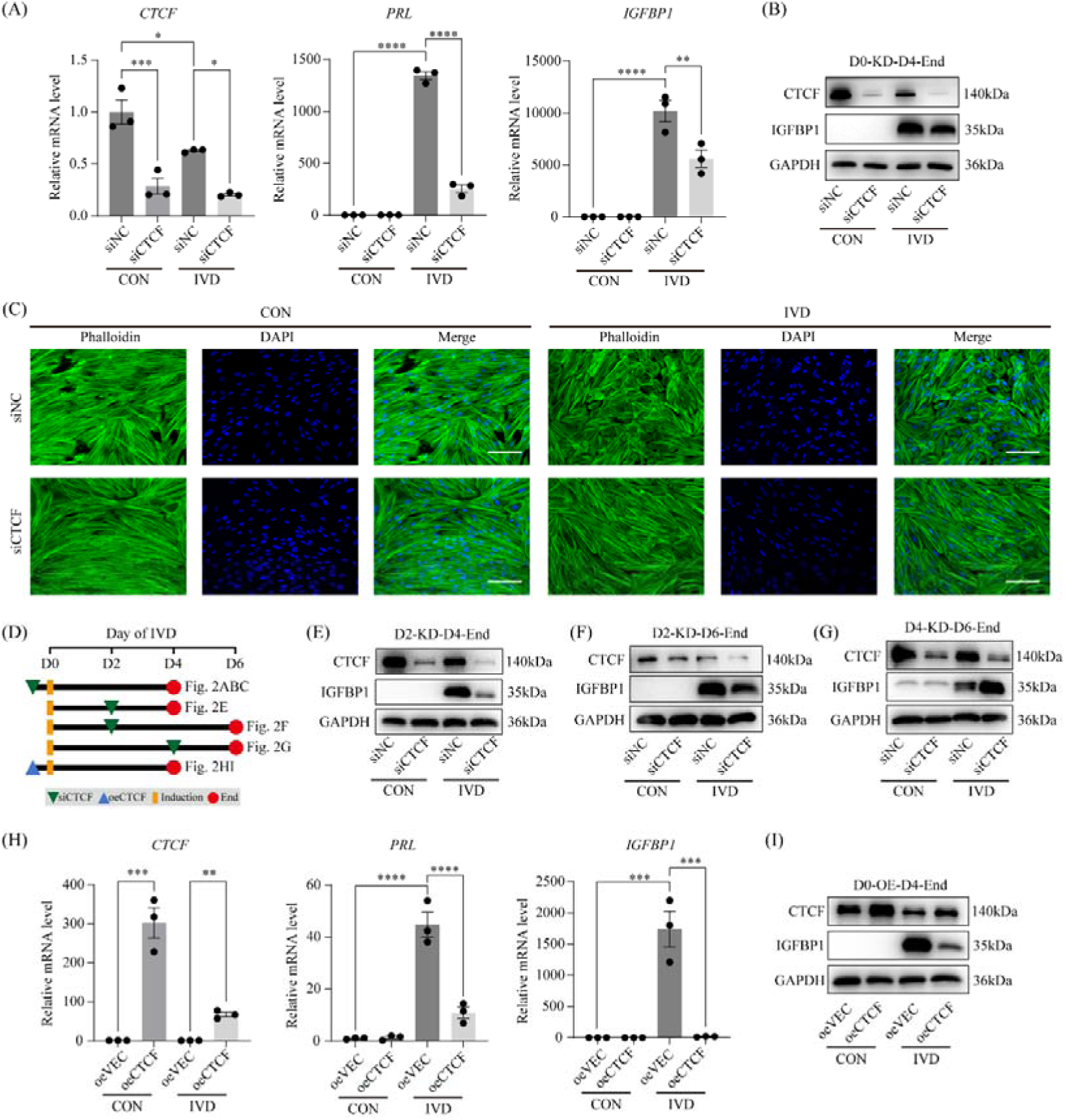
CTCF is essential for the early but dispensable for the late stage of decidualization. (A) Quantitative RT-PCR measures *PRL* and *IGFBP1* RNA levels in hESCs after CTCF knockdown and 4-day decidualization. siNC, non-targeting control; siCTCF, CTCF-targeting siRNA. Data are presented as mean ± SEM. *, P < 0.05; **, P < 0.01; ***, P < 0.001; ****, P < 0.0001. (B) IGFBP1 protein expression after CTCF knockdown and 4-day decidualization. (C) Phalloidin staining of F-actin in decidualized hESCs after siRNA-mediated CTCF knockdown. Bar=100 μm. (D) Diagram illustrates knockdown and overexpression time points to elucidate CTCF’s role in decidualization. (E) IGFBP1 protein levels in decidualized hESCs at day 4, transfected with CTCF siRNA at day 2 post-decidualization induction. (F) IGFBP1 protein levels in decidualized hESCs at day 6, transfected with CTCF siRNA at day 2 post-decidualization induction. (G) IGFBP1 protein levels in decidualized hESCs at day 6, transfected with CTCF siRNA at day 4 post-decidualization induction. (H) Quantitative RT-PCR assesses *PRL* and *IGFBP1* mRNA expression in hESCs after CTCF overexpression and 4-day decidualization. oeVEC, empty vector control; oeCTCF, CTCF overexpression vector. Data are presented as means ± SEM. **, P < 0.01; ***, P < 0.001; ****, P < 0.0001. (I) IGFBP1 protein expression after CTCF overexpression and 4-day decidualization.

To ascertain the evolutionary conservation of CTCF’s role in decidualization, we performed siRNA-mediated knockdown in primary mouse endometrial stromal cells (mESCs) before in vitro decidualization induction. Notably, CTCF depletion significantly reduced *Prl8a2* expression, a canonical marker of murine decidualization (**Fig. S5A-B**). These findings confirm the functional conservation of CTCF as a critical regulator of decidualization across humans and mice.

### CTCF primes hESCs for decidualization via HOXA11

To uncover genes regulated by CTCF in hESCs, we performed RNA-seq analysis on mRNAs isolated from CTCF-knockdown hESCs and control cells. Data analysis revealed 424 differentially expressed genes, including 195 up-regulated genes and 229 down-regulated genes (**Fig. 3A** and **Table S1**). Gene Ontology (GO) analysis demonstrated significant enrichment of downregulated genes in pathways regulating cell proliferation and migration (**Fig. 3B**), consistent with our functional studies (**Fig. S4D-E** and **G**). Notably, RNA-seq analysis revealed HOXA11, which is a master regulator of decidualization, as among the significantly downregulated genes in CTCF-depleted hESCs (**Fig. 3C**). Subsequent validation through quantitative RT-PCR and western blot confirmed HOXA11 suppression following CTCF siRNA treatment (**Fig. 3D-E**) and markedly upregulated HOXA11 expression with CTCF overexpression (**Fig. 3F-G**). To ascertain whether the CTCF-HOXA11 axis is conserved in mice, we conducted siRNA-mediated knockdown of CTCF in isolated mESCs. Knockdown of CTCF in mESCs similarly resulted in a significant inhibition of HOXA11 expression **(Fig. S6A-B**). These findings uncovered HOXA11 as a putative target gene of CTCF.

**Figure 3.**
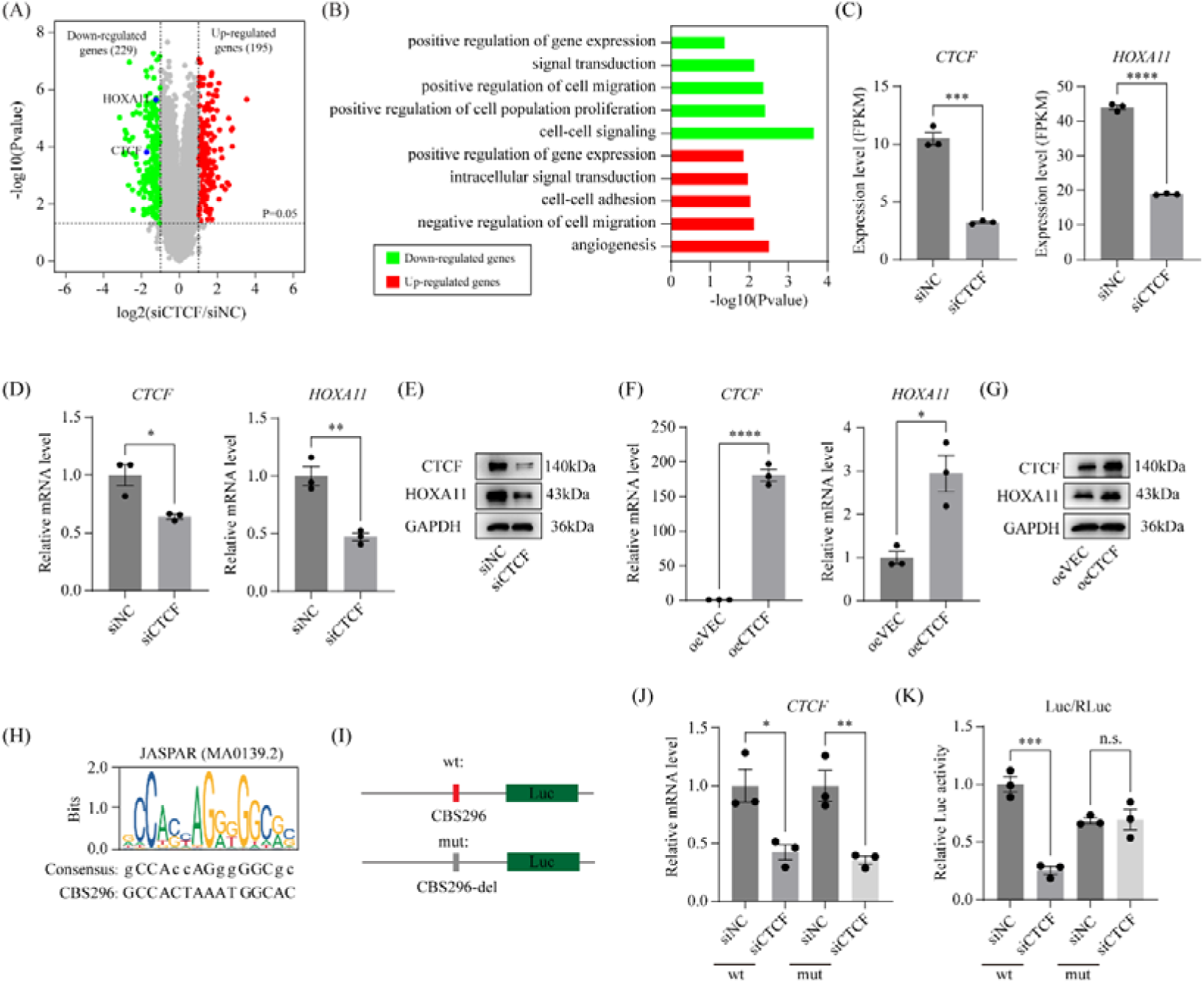
HOXA11 serves as a direct target of CTCF. (A) A volcano plot depicts significantly expressed genes in hESCs following 48h CTCF knockdown, as identified by RNA-seq (fold change > 2, P < 0.05). (B) Gene ontology enrichment analysis of differentially expressed genes. (C) RNA-seq-derived HOXA11 expression levels in hESCs after CTCF knockdown. Data are presented as mean ± SEM. ***, P < 0.001; ****, P < 0.0001. (D-E) Quantitative RT-PCR (D) and western blot (E) validate HOXA11 expression in hESCs after 48h CTCF knockdown. Data are presented as mean ± SEM. *, P < 0.05; **, P < 0.01. (F-G) HOXA11 mRNA (F) and protein (G) levels in hESCs following 48h CTCF overexpression. Data are presented as mean ± SEM. *, P < 0.05; ****, P < 0.0001. (H) Sequence of the CTCF binding site CBS296 in the HOXA11 promoter. (I) Schematic diagram of the luciferase reporter vector, incorporating the HOXA11 promoter (from −500 bp to 500 bp of TSS) upstream of the luciferase gene in pGL3-basic. (J-K) Impact of CBS296 on luciferase reporter gene expression in HEK293T cells co-transfected with the reporter vector and either siCTCF or siNC for 48h. Renilla luciferase in pRL-TK vector serves as normalization control. Data are presented as mean ± SEM. *, P < 0.05; **, P < 0.01; ***, P < 0.001.

Subsequently, we investigated the regulatory mechanism how HOXA11 is regulated by CTCF. As fundamental units of higher-order chromatin architecture, topologically associating domains (TADs) compartmentalize DNA functional elements such as enhancers and insulators, which coordinate spatial regulation of neighboring genes [25, 26]. To identify regulatory elements for HOXA11, we analyzed mouse uterine Hi-C data [27] and observed the HOXA cluster 3’-end localized at a TAD boundary, while its 5’-end resided within the TAD interior (**Fig. S7A**). Additionally, mouse uterine CTCF ChIP-seq data [23] showed 5 CTCF binding sites (CBSs) within the HoxA cluster: at the intergenic region between HOXA5 and HOXA6 (CBS5|6), between HOXA6 and HOXA7 (CBS6|7), between HOXA7 and HOXA9 (CBS7|9), between HOXA10 and HOXA11 (CBS10|11), and between HOXA11 and HOXA13 (CBS11|13) (**Fig. S7B**). Previous studies have shown that these sites regulate the expression of posterior HOXA9-HOXA13 genes [28, 29]. Notably, we identified a novel CTCF binding site, CBS296, near the HOXA11 TSS that had not been previously characterized (**Fig. S7C** and **Fig. 3H**). Functional assessment using luciferase reporters containing wild-type or CBS296-deleted *HOXA11* promoter segments demonstrated that CTCF knockdown specifically reduced reporter activity when CBS296 was intact, but not following its deletion (**Fig. 3I-K**). These findings establish CBS296 as a critical cis-element mediating CTCF-dependent regulation of HOXA11.

Leveraging the publicly accessible microarray dataset (GSE4888), we observed a gradual decline in HOXA11 mRNA levels during the secretory phase of the menstrual cycle (**Fig. 4A**). In hESCs, our study revealed that HOXA11 expression peaks on day 2 and subsequently declines during in vitro decidualization (**Fig. 4B-C**). Interestingly, HOXA11 knockdown only on day 1 impeded decidualization, whereas knockdown at other timepoints, including days 0, 2, 3, and 4, promoted decidualization (**Fig. 4D-L**). Furthermore, overexpressing HOXA11 on day 0 impaired decidualization, whereas overexpressing it on day 1 enhanced decidualization (**Fig. 4M-N**). These observations indicate that HOXA11 is exclusively required on day 1 for decidualization. Consequently, we found that overexpressing HOXA11 on day 1, but not on day 0, effectively rescued decidualization in hESCs with CTCF knockdown (**Fig. 4O-R**). These findings suggest that HOXA11 acts as a pivotal downstream effector of CTCF in hESCs during the decidualization process.

**Figure 4.**
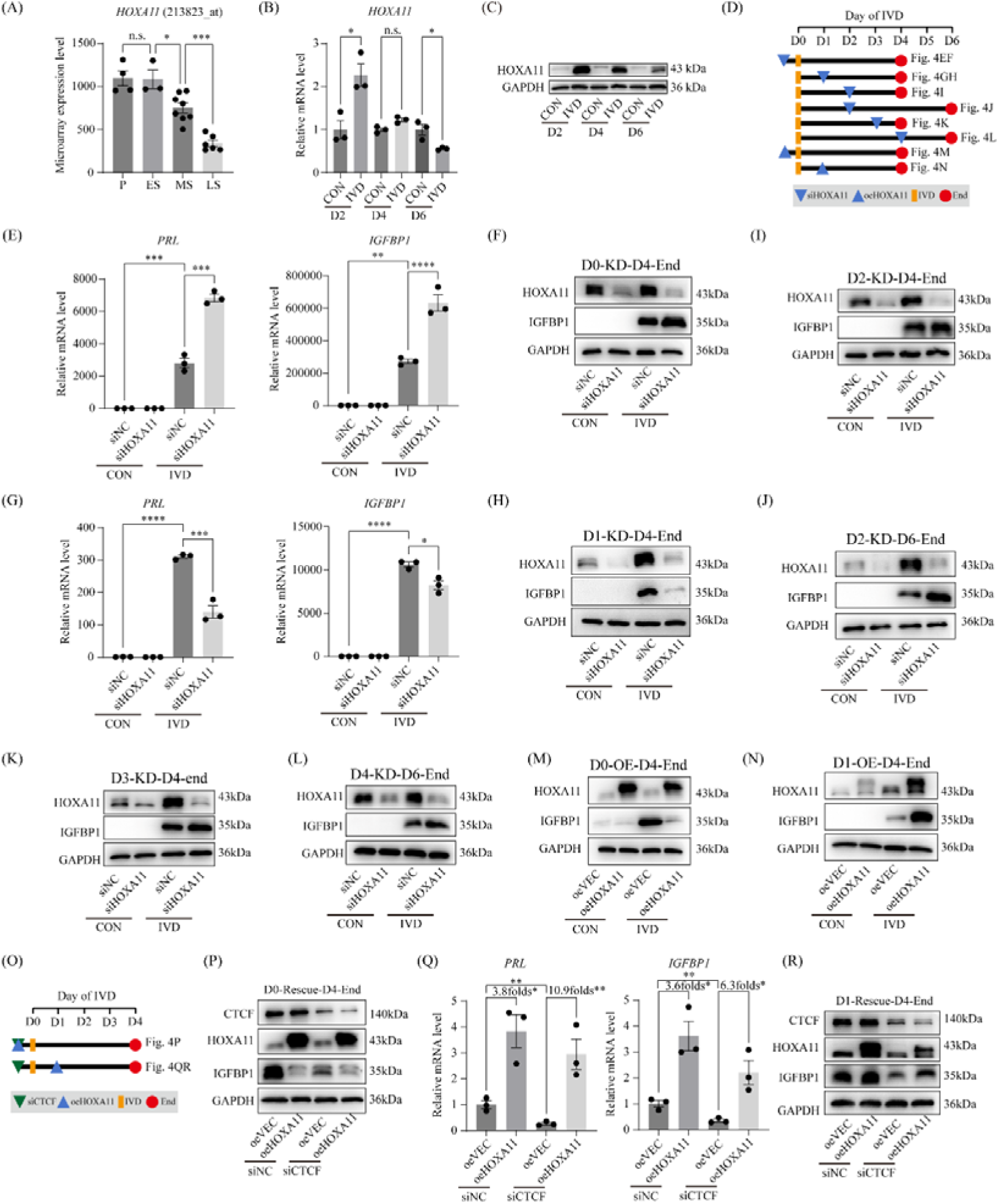
HOXA11 overexpression rescues decidualization impairment induced by CTCF knockdown. (A) Microarray dataset GSE4888 reveals *HOXA11* mRNA expression in human endometrium across the menstrual cycle. P, proliferative phase; ES, early secretory phase; MS, middle secretory phase; LS, late secretory phase. Data are presented as means ± SEM. n.s., not significant. *, P < 0.05; ***, P < 0.001. (B-C) HOXA11 mRNA (B) and protein (C) expression in decidualizing hESCs at different time points. CON, vehicle control; IVD, in vitro decidualization. Data are presented as mean ± SEM. *, P < 0.05. (D) Diagram illustrates knockdown and overexpression time points to elucidate HOXA11’s role in decidualization. (E-N) PRL and IGFBP1 expression levels in decidualized hESCs after HOXA11 knockdown or overexpression at various time points. Data are presented as mean ± SEM. *, P < 0.05; **, P < 0.01; ***, P < 0.001; ****, P < 0.0001. (O) Diagram depicts HOXA11 overexpression to rescue decidualization impairment following CTCF knockdown. (P-R) PRL and IGFBP1 expression levels in decidualized hESCs with CTCF knockdown and rescued by HOXA11 overexpression. Data are presented as mean ± SEM. *, P < 0.05; **, P < 0.01.

### *Pgr*-Cre-mediated deletion of CTCF in female mice causes infertility due to embryo implantation failure

To elucidate CTCF’s role in the mouse uterus, we generated conditional *Ctcf* knockout mice (*Ctcf*^d/d^) by crossing Ctcf-floxed mice (*Ctcf*^f/f^) with mice expressing Cre recombinase under the *Pgr* promoter (*Pgr*^Cre/+^) (**Fig. 5A-B** and **Fig. S8A-C**). PCR analysis confirmed successful excision of exon 8, flanked by floxp sites, by Cre recombinase in the uteri of adult female mice (**Fig. 5C**). To assess the impact of Ctcf deletion on female fertility, we conducted a 6-month fertility test by pairing *Ctcf*^d/d^ and *Ctcf*^f/f^ adult female mice with fertile wild-type male mice of the same genetic strain. Our results revealed that *Ctcf*^d/d^ female mice were infertile, whereas *Ctcf*^f/f^ control female mice maintained normal fertility (**Fig. 5D**), underscoring CTCF’s indispensable role in achieving successful pregnancy.

**Figure 5.**
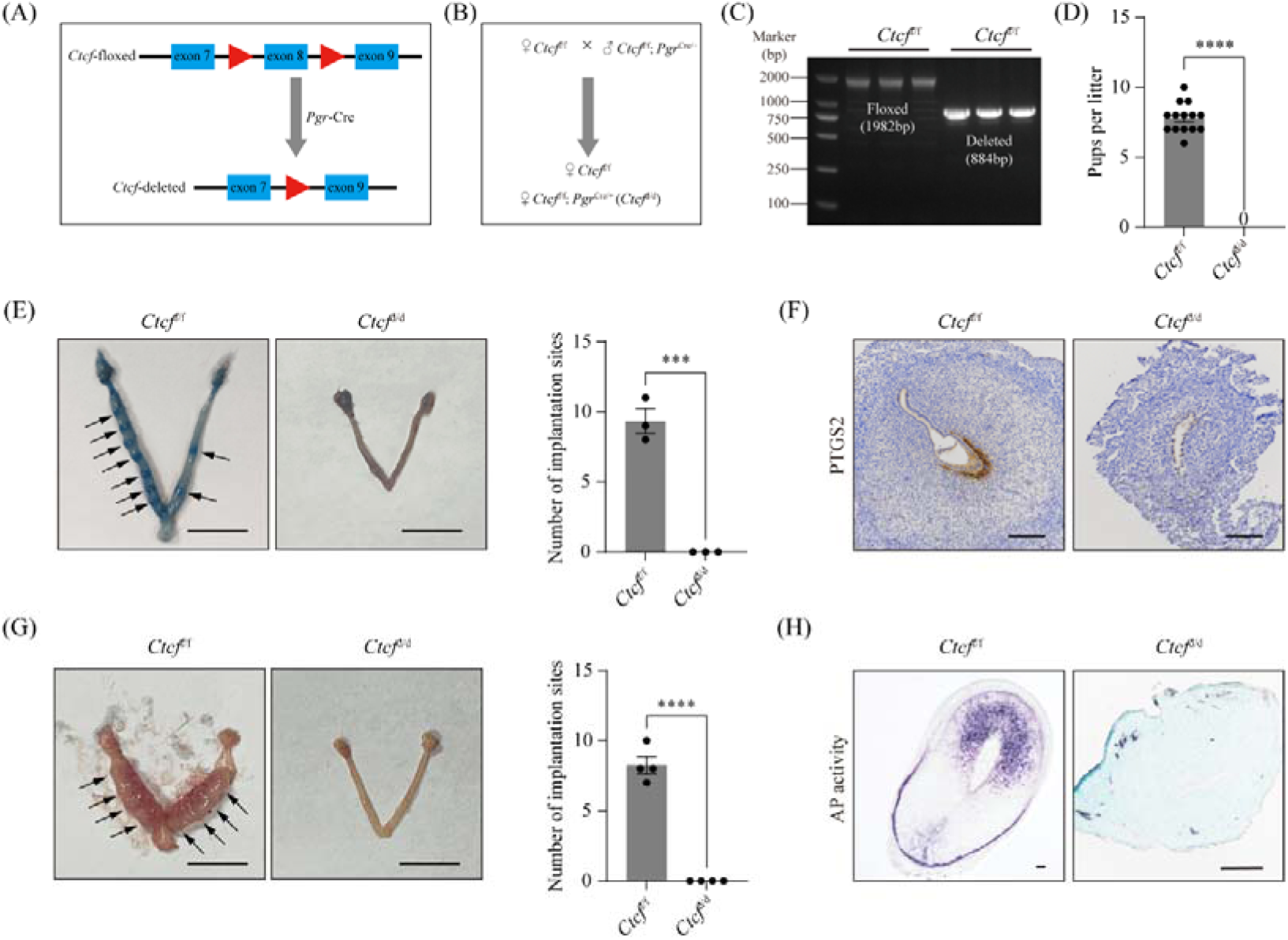
*Pgr*-Cre-mediated deletion of CTCF in mice leads to female infertility attributed to implantation failure. (A) Diagram outlines the strategy for conditional CTCF deletion. *Ctcf*^f/f^ mice, carrying two loxP sites flanking exon 8, were crossed with *Pgr*-Cre driver line to generate *Ctcf*^d/d^ mice. (B) Diagram of the breeding scheme. (C) PCR analysis confirms knockout efficiency of *Ctcf*^d/d^ mice at the DNA level in the uterus at 8 weeks of age. (D) Comparison of litter sizes between 3 *Ctcf*^d/d^ and 3 *Ctcf*^f/f^ mice over a 6-month period. Data are presented as mean ± SEM. ****, P < 0.0001. (E) Bar plot illustrates the number of embryo implantation sites in *Ctcf*^d/d^ and *Ctcf*^f/f^ mice on GD5. Arrowheads mark implantation sites, which are visualized by blue dye injection. Bar=1 cm. Data are presented as mean ± SEM. ***, P < 0.001. (F) Immunohistochemistry staining of PTGS2 at implantation sites in *Ctcf*^d/d^ and *Ctcf*^f/f^ uteri on GD5. Bar=100 μm. (G) Bar plot shows the number of embryo implantation sites in *Ctcf*^d/d^ and *Ctcf*^f/f^ mice on GD8. Bar=1 cm. Data are presented as mean ± SEM. ****, P < 0.0001. (H) Alkaline phosphatase staining of the implantation site from *Ctcf*^d/d^ and *Ctcf*^f/f^ mice on GD8. Bar=100 μm.

To identify the cause of infertility in *Ctcf*^d/d^ mice, we examined their pregnancy status at different timepoints. Notably, on GD5 and GD8, *Ctcf*^d/d^ mice exhibited a complete absence of implantation sites (**Fig. 5E-H**). We therefore conclude that CTCF is essential for normal embryo implantation.

### Loss of CTCF results in impaired receptivity

We subsequently explored whether implantation failure in *Ctcf*^d/d^ females stemmed from ovarian dysfunction. Immunohistochemical analysis revealed that CTCF expression remained unaltered in the ovaries of *Ctcf*^d/d^ mice on GD4 (**Fig. S9A**). Furthermore, critical steroid biosynthetic enzymes, CYP11A1 (P450scc) and HSD3B2, were normally expressed in the corpus luteum of these mice (**Fig. S9B**). Additionally, serum P_4_ levels were comparable between *Ctcf*^d/d^ and control *Ctcf*^f/f^ female mice on GD4 (**Fig. S9C**), indicating normal ovarian activity.

We further examined uterine receptivity. On GD4, the uterine size and weight of *Ctcf*^d/d^ mice were significantly reduced compared to those of *Ctcf*^f/f^ mice (**Fig. 6A**). Quantitative RT-PCR data revealed that *Ctcf* knockout was associated with down-regulation of *Ctcf* mRNA, potentially due to nonsense-mediated mRNA decay (NMD) (**Fig. 6B**). The loss of CTCF at the protein level was confirmed via western blot analysis (**Fig. 6C**). Immunohistochemical analysis demonstrated efficient deletion of CTCF in stromal cells but preservation in luminal and glandular epithelial cells (**Fig. 5D**), consistent with previous findings [23]. Histological analysis revealed a reduced stroma layer and a decrease in the number of endometrial glands in the *Ctcf*^d/d^ uterus (**Fig. 5E**). Next, we evaluated uterine receptivity markers. The loss of the anti-adhesive glycoprotein mucin 1 (MUC1) on the entire luminal epithelium serves as an indicator of uterine receptivity in mice. Immunohistochemical analysis showed abundant MUC1 protein on the luminal epithelium of *Ctcf*^d/d^ mice, whereas it was absent on the luminal epithelium of *Ctcf*^f/f^ mice. The cessation of luminal epithelium proliferation and the initiation of stromal cell proliferation are pivotal for establishing uterine receptivity. Immunohistochemical staining for MKI67, a marker of proliferative cells, indicated that although luminal epithelium proliferation ceased in *Ctcf*^d/d^ mice on GD4, stromal cell proliferation was absent (**Fig. 5F**). RNA-seq analysis identified 1422 differentially expressed genes, with 673 upregulated and 749 downregulated in *Ctcf*^d/d^ mice relative to *Ctcf*^f/f^ mouse uterus (**Fig. S10A**). RNA-seq data revealed that *Ctcf* deletion was linked to decreased expression of *Hoxa11*, while the expression of *Pgr* and *Esr1* remained unchanged (**Fig. S10B-D**). These findings were further corroborated by quantitative RT-PCR (**Fig. 6B**), western blot (**Fig. 6C**) and immunohistochemical analyses (**Fig. 6D**). Collectively, these results suggest that the deletion of CTCF in uterine stromal cells disrupts uterine receptivity by reducing HOXA11 expression.

**Figure 6.**
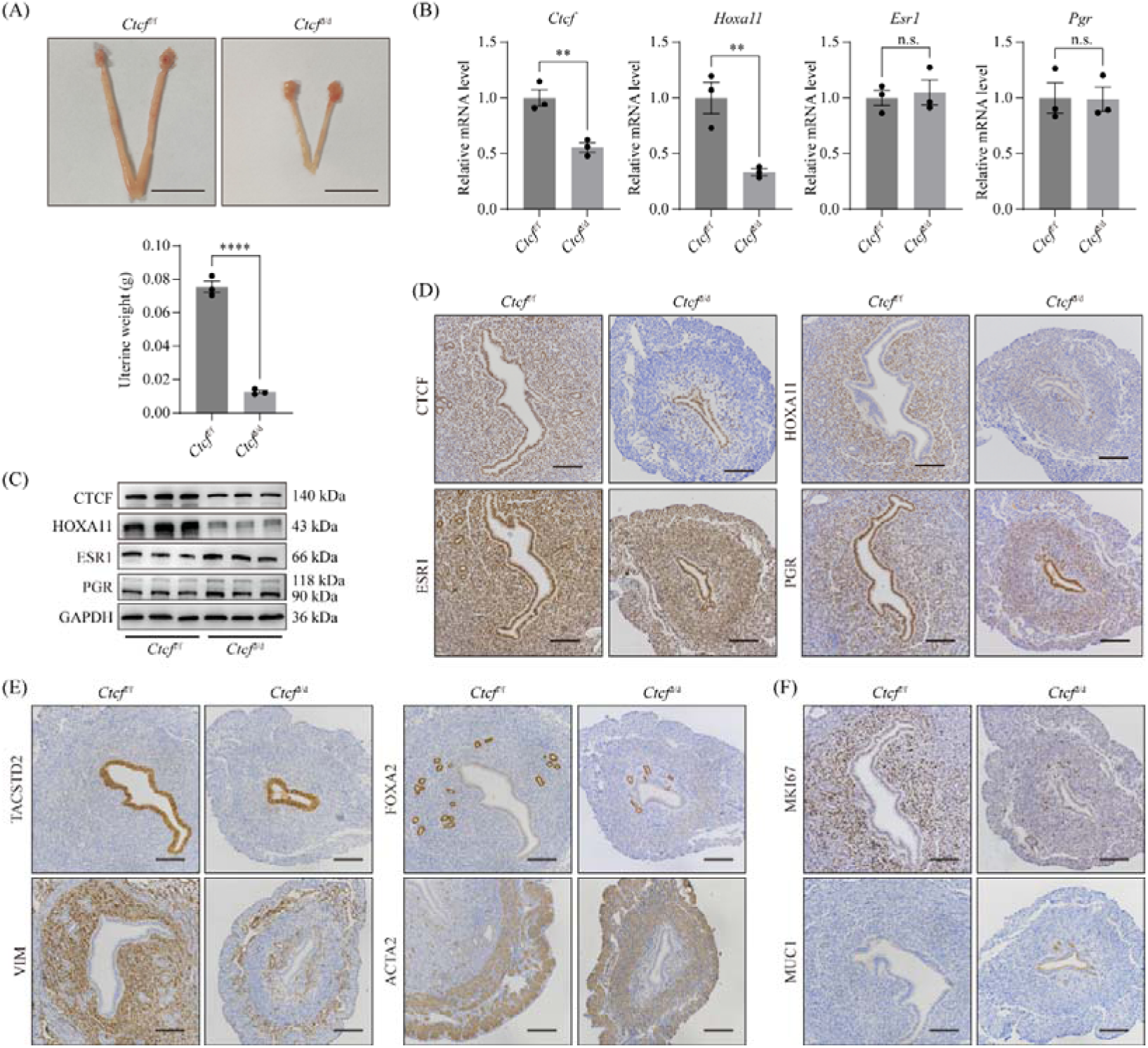
Loss of CTCF compromises uterine receptivity. (A) Bar plot depicts uterine weight in *Ctcf*^d/d^ and *Ctcf*^f/f^ mice on GD4. Bar=1 cm. Data are presented as mean ± SEM. ****, P < 0.0001. (B) Quantitative RT-PCR analysis of *Ctcf*, *Hoxa11*, *Esr1*, and *Pgr* expression in *Ctcf*^d/d^ and *Ctcf*^f/f^ uteri on GD4. Data are presented as means ± SEM. **, P < 0.01. (C) Western blot analysis of CTCF, HOXA11, ESR1, and PGR protein levels in the uterus of *Ctcf*^d/d^ and *Ctcf*^f/f^ mice on GD4. (D-F) Immunohistochemistry analysis of marker genes in the uterus of *Ctcf*^d/d^ and *Ctcf*^f/f^ mice on GD4. Bar=100 μm.

### Loss of CTCF leads to decidual response defect

Drawing from insights gained through hESC studies, we hypothesized that *Ctcf* deletion would impair uterine decidualization during murine pregnancy. To investigate this, we employed an artificial decidualization model. Remarkably, while *Ctcf*^f/f^ uteri demonstrated robust decidualization, *Ctcf*^d/d^ counterparts failed to exhibit this response (**Fig. 7A**). In line with defective stromal differentiation, alkaline phosphatase activity, a characteristic marker of decidualization, was evident in *Ctcf*^f/f^ uteri but absent in *Ctcf*^d/d^ uteri (**Fig. 7B**). Considering that decidualizing stromal cells secrete pro-angiogenic factors, we evaluated vascular development through PECAM1 immunostaining. As anticipated, *Ctcf*^d/d^ uteri exhibited significantly reduced endothelial cell staining compared to *Ctcf*^f/f^ controls (**Fig. 7C**). Quantitative RT-PCR analysis of canonical decidualization markers (*Bmp2*, *Wnt4*, *Prl8a2*) further substantiated the functional impairment in *Ctcf*^d/d^ uteri (**Fig. 7D**). Notably, *Ctcf* ablation was associated with decreased HOXA11 expression in uterine stromal cells (**Fig. 7E-F**). Collectively, these findings establish CTCF as a crucial regulator of uterine decidualization in mice.

**Figure 7.**
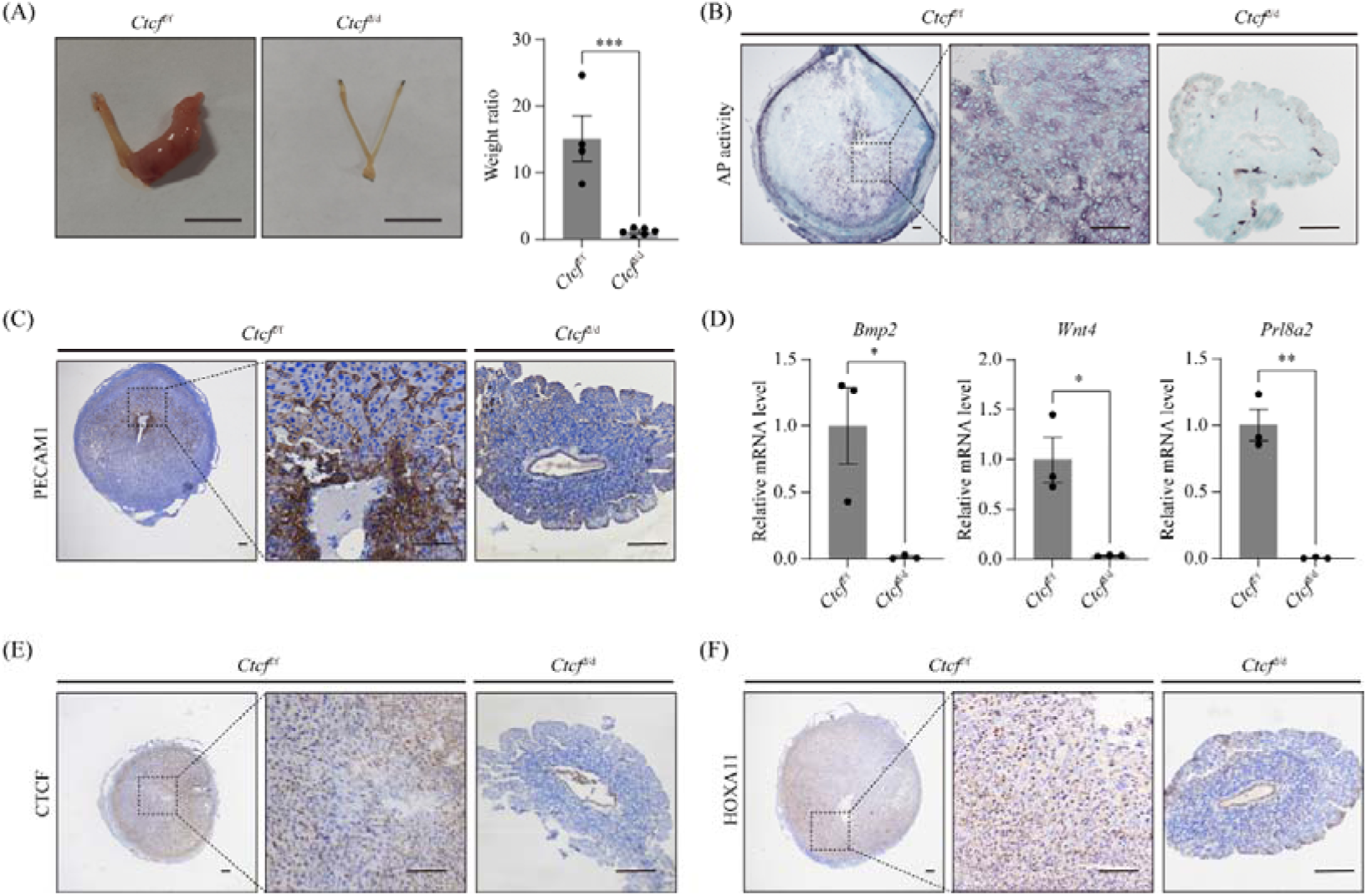
Loss of CTCF induces a decidual response defect. (A) Bar plot illustrates the weight ratio of the stimulated to unstimulated uterine horn in *Ctcf*^d/d^ and *Ctcf*^f/f^ mice post-artificial decidualization. Bar=1 cm. Data are presented as mean ± SEM. ***, P < 0.001. (B) Alkaline phosphatase (AP) staining of the stimulated uterine horn in *Ctcf*^d/d^ and *Ctcf*^f/f^ mice. Bar=100 μm. (C) Immunohistochemistry staining of PECAM1 in the stimulated uterine horn of *Ctcf*^d/d^ and *Ctcf*^f/f^ mice. Bar=100 μm. (D) Quantitative RT-PCR analysis of decidualization marker genes in the stimulated uterine horn of *Ctcf*^d/d^ and *Ctcf*^f/f^ mice following artificial decidualization. Data are presented as means ± SEM. *, P < 0.05; **, P < 0.01. (E-F) Immunohistochemistry staining of CTCF and HOXA11 in the stimulated uterine horn of *Ctcf*^d/d^ and *Ctcf*^f/f^ mice post-artificial decidualization. Bar=100 μm.

## Discussion

CTCF is a versatile transcriptional regulator and chromatin architectural protein required for mammalian development. Here, we demonstrate that CTCF depletion in human endometrial stromal cells (hESCs) impairs decidualization competence, with HOXA11 downregulation identified as a critical mediator of this defect. Mechanistic investigations revealed direct CTCF binding to the HOXA11 promoter region, facilitating its transcriptional activation. In murine models, *Pgr*-Cre-mediated uterine stromal CTCF ablation resulted in complete implantation failure attributable to dual defects in uterine receptivity acquisition and decidualization capacity. This study elucidates the physiological role of CTCF in the uterus during pregnancy.

In this study, we employed both in vitro and in vivo models to dissect the role of CTCF in decidualization. In human endometrial stromal cells (hESCs), CTCF knockdown at decidualization initiation (day 0) or early stage (day 2) significantly impaired differentiation capacity, whereas late-stage knockdown (day 4) unexpectedly enhanced this process. Conversely, forced CTCF overexpression disrupted proper decidualization progression. Given that CTCF expression is down-regulated during the decidualization process, our findings suggest that CTCF expression in hESCs is essential for the early phase of decidualization, while excessive CTCF expression in the late stage of decidualization is likely harmful. In *Pgr*-Cre-mediated CTCF conditional knockout mice, we observed that the uterine deletion of CTCF unusually occurred only in stromal cells but not in epithelial cells. This is consistent with a previous study, which showed that while uterine CTCF deletion initially causes loss of epithelial CTCF expression by postnatal day 21 (PND21), re-emergence occurs during pubertal development, likely through selective proliferation of CTCF-positive epithelial clones [23]. The *Ctcf*^d/d^ uteri had a reduced stroma layer and a decrease in the number of endometrial glands, which can be traced back to pubertal uterine development as described previously [23]. Functionally, *Ctcf*^d/d^ uteri display defective uterine receptivity and compromised decidualization. Taken together, our in vitro and in vivo data consistently showed that CTCF is required for decidualization, especially the initial stage of decidualization.

In this study, we identified HOXA11 as a critical downstream effector of CTCF in the uterus. In hESCs, CTCF depletion reduces HOXA11 expression while CTCF overexpression enhances it. Notably, during in vitro decidualization, CTCF expression peaks at day 0, whereas HOXA11 peaks at day 2, with a 2-day lag. Functionally, HOXA11 overexpression rescues decidualization defects caused by CTCF knockdown, establishing HOXA11 as a key mediator of CTCF’s effect in decidualization. In *Pgr*-Cre-mediated CTCF conditional knockout mice, stromal CTCF deletion phenocopies HOXA11 knockout, including reduced stromal compartment thickness, hypoplastic endometrial glands, impaired uterine receptivity, and defective decidualization [30, 31]. Mechanistically, we identified a novel CTCF binding site within the HOXA11 promoter that directly promotes its transcription. Collectively, this study demonstrated that CTCF safeguards HOXA11 expression in the uterus. Our findings hold significant clinical importance, as reduced HOXA11 expression is often associated with uterine pathophysiology, such as endometriosis [32–34], spontaneous pregnancy loss [34], and chronic endometritis [35].

In summary, this study elucidates a conserved regulatory axis wherein CTCF orchestrates uterine decidualization critical for pregnancy establishment through HOXA11-dependent mechanisms.

## Materials and Methods

### Human endometrial sample collection

Participants for this study were recruited from the Reproductive Medicine Center at the First Affiliated Hospital of Sun Yat-sen University in China from July 2020 to October 2021. All participants provided informed consent, and the study received approval from the Ethical Committee of the First Affiliated Hospital of Sun Yat-sen University (Approval No. 2018-266). We selected normally fertile participants who exhibited no evident endometrial pathology and achieved a confirmed clinical pregnancy following embryo transfer. Comprehensive patient details are outlined in our prior study [36].

### Isolation and culture of human endometrial stromal cells (hESCs)

Three healthy, fertile participants with no evidence of endometrial pathology and a clinically confirmed pregnancy following embryo transfer were recruited for this study. The protocol received prior approval from the Ethical Committee of the First Affiliated Hospital of Sun Yat-sen University (Approval No. 2018-266), and all participants provided written informed consent. A summary of the patients’ characteristics is presented in **Table S3**. Endometrial tissues were dissected into fragments and digested with type I collagenase (Gibco) for 1 hour. Endometrial epithelial and stromal cells were then separated using membrane filters (100 µm and 40 µm cell filters, Corning). The hESCs were cultured in DMEM/F12 medium (Gibco) supplemented with 10% charcoal-stripped fetal bovine serum (cFBS, VivaCell). For decidualization induction, cells were treated with 0.5 mM 8Br-cAMP (Sigma) and 1 μM medroxyprogesterone acetate (Sigma) in 2% cFBS.

### Mice

The *Ctcf*^f/f^ mice (Cat. No. NM-CKO-200031) and *Pgr*^Cre/+^ mice (Cat. No. NM-KI-200117) were obtained from Shanghai Model Organisms Center, Inc., China. The *Ctcf*^d/d^ mice were generated by crossing *Ctcf*^f/f^ mice with *Pgr*^Cre/+^ mice, with *Ctcf*^f/f^ littermates serving as the wild-type control group. Adult female mice were mated with either fertile males to induce pregnancy or vasectomized males to induce pseudo-pregnancy. The day of vaginal plug detection was designated as gestational day 1 (GD1). For artificial decidualization experiments, a 20 μl volume of sesame oil (Sigma) was injected into one uterine horn of pseudo-pregnant mice on GD4, and uterine samples were collected on GD8. All animal procedures were approved by the Institutional Animal Care and Use Committee of South China Agricultural University (Approval No. 2022B042).

### Isolation and culture of mouse endometrial stromal cells (mESCs)

The isolation and culture of mouse endometrial stromal cells (mESCs) were conducted with minor modifications to a previously established protocol [37]. Briefly, uterine horns from adult mice on gestational day 4 (GD4) were dissected into small fragments. These fragments were submerged in Hank’s balanced salt solution (HBSS) containing 6 mg/ml dispase and 25 mg/ml trypsin for 1 hour at 4°C. Subsequently, the tissues were incubated for an additional hour at room temperature, followed by 10 minutes at 37°C. After removing endometrial epithelial clumps, the remaining tissues were incubated once more in HBSS containing 0.5 mg/ml collagenase at 37°C for 30 minutes. To obtain stromal cells, the digested cell suspension was filtered through a 70-μm mesh. The isolated cells were seeded onto 60-mm dishes at a density of 5 × 10L cells per dish and cultured in a 1:1 mixture of phenol red-free Dulbecco’s modified Eagle’s medium and Ham F-12 nutrient medium (DMEM/F12; Gibco), supplemented with 10% charcoal-stripped fetal bovine serum (cFBS; Biological Industries) and antibiotics. Following an initial 2-hour incubation, the medium was replaced with phenol red-free DMEM/F12 containing 1% cFBS, 10 nM estradiol (E_2_), and 1 μM progesterone (P_4_) to induce decidualization.

### Quantitative RT-PCR

Total RNA was extracted using TRIzol reagent (Invitrogen). Subsequently, 1 μg of the purified RNA was employed to synthesize cDNA with HiScript III RT Super Mix (Vazyme). Quantitative PCR was performed using ChamQ SYBR qPCR Master Mix (Vazyme) on an Applied Biosystems 7300 Plus (Life Technologies). Expression data were normalized against *Rpl7* as the reference gene. A comprehensive list of the primer sequences used in this study is provided in **Table S4**.

### Western blot

Uterine tissues were homogenized, and cultured cells were lysed using Radio-Immunoprecipitation Assay (RIPA) buffer supplemented with protease inhibitors (Roche). Total protein extracts were resolved on a 10% SDS-PAGE gel and transferred to a PVDF membrane (Millipore). Before antibody probing, the membrane was blocked with 5% skim milk in Tris-buffered saline with Tween 20 (TBST). The membrane was then incubated overnight at 4°C with the primary antibody. Following thorough washing, it was incubated with a horseradish peroxidase (HRP)-conjugated secondary antibody for 1 hour at room temperature. Immunoreactive signals were visualized using the ECL chemiluminescent kit (Amersham Biosciences). GAPDH bands served as the loading control. The primary antibodies used in this study are listed in **Table S5**.

### Immunofluorescence

Cellular fixation was carried out using 4% paraformaldehyde (P6148, Sigma) for 15 minutes, followed by permeabilization with 0.1% Triton X-100 (T8787, Sigma) for 10 minutes. Non-specific binding was minimized by blocking with 10% goat serum at 37°C for 1 hour. A primary antibody targeting the protein of interest was applied and incubated overnight at 4°C. The next day, samples were incubated with species-specific secondary antibodies (Jackson ImmunoResearch) for 40 minutes at room temperature. Immunofluorescence staining with phalloidin was performed as per the manufacturer’s instructions (Beyotime, C2201S). The primary antibodies used in this study are listed in **Table S5**.

### RNA-seq

Uterine segments were rapidly frozen in liquid nitrogen and stored at −80 °C until further processing. Total RNA extraction, cDNA library construction, and high-throughput DNA sequencing were conducted according to our previously established protocols [38]. Following quality control, the sequence data were aligned to the mouse genome (UCSC mm10) using HISAT v2.2.1 [39]. Gene-level expression values were calculated from the aligned sequences with Cufflinks v2.2.1 [40]. Differentially expressed genes were identified based on a fold change > 2 and an adjusted P-value < 0.05. To elucidate the functional roles of these differentially expressed genes, gene ontology analysis was performed using the DAVID online tools with knowledgebase v2024q4 [41], and redundant enriched terms were manually removed.

### Immunohistochemistry

Formalin-fixed, paraffin-embedded uterine tissues were sectioned into 5-μm slices. Heat-induced antigen retrieval was performed in a 10 mM citrate buffer (pH = 6.0) for 15 minutes. To quench endogenous peroxidase activity, sections were treated with 3% HLOL for 20 minutes. After blocking with 10% horse serum in PBS, sections were incubated with the primary antibody overnight at 4°C. Subsequently, they were incubated with an HRP-conjugated secondary antibody for one hour at room temperature. Immunoreactive signals were visualized using a DAB staining kit (Zhongshan Golden Bridge Biotechnology Co., Beijing). For contrast, sections were counterstained with hematoxylin. The primary antibodies used in this study are listed in **Table S5**.

### Alkaline phosphatase staining

Frozen tissue sections, precisely cut at 10 μm, were rapidly fixed in cold acetone for 15 minutes and then thoroughly washed three times with PBS. Alkaline phosphatase activity was visualized using the BCIP/NBT kit (Zhongshan Golden Bridge Biotechnology Co., Beijing), adhering to the manufacturer’s recommended protocol. For counterstaining, a 1% methyl green solution was applied.

### Luciferase reporter assay

The genomic segment of the human HOXA11 promoter (spanning from −500bp to +500bp relative to the transcription start site) was synthesized and cloned into the pGL3-basic vector (Promega). Site-directed mutagenesis was employed to generate the CBS147-deletion mutant. The reconstructed plasmid and siRNA were co-transfected into HEK293T cells using Lipofectamine 3000 (Invitrogen). *Renilla* luciferase from the pRL-TK vector served as the normalization control. Cell lysates were harvested 48 hours post-transfection. Luciferase activity was assessed using the Dual-Luciferase Reporter Assay System (Promega), with firefly luciferase activity normalized to *Renilla* luciferase activity. The detailed information on the siRNAs used is provided in **Table S6**.

### Statistical analysis

Statistical analyses were conducted using GraphPad Prism v10.0.0. The two-tailed unpaired Student’s t-test was employed to compare the means of two groups, while the one-way ANOVA with Tukey’s multiple comparison test was used for comparing the means of more than two groups. Data are expressed as the mean ± SEM, and statistical significance was defined as P < 0.05.

## Acknowledgements

This research was funded by National Natural Science Foundation of China (32370913 and 32070845), Guangdong Natural Science Funds for Distinguished Young Scholars (2021B1515020079), Innovation Team Project of Guangdong University (2019KCXTD001), Guangdong Special Support Program (2019BT02Y276), National Key R&D Program of China (2018YFA0801404), and Double first-class discipline promotion project (2023B10564003).

## Author contributions

Y.-W.X., S.-H.Y. and J.-L.L. designed research; X.-Q.Y., R.-F.J., A.-N.Z., Y.-M.D., S.T., and Q.-Y.Z. performed the experiments; X.-Q.Y., Y.-W.X., S.-H.Y. and J.-L.L. analyzed the data. X.-Q.Y., Y.-W.X., S.-H.Y. and J.-L.L. wrote the paper. All authors read and approved the final paper.

## Data availability statements

The sequencing data generated in this study are deposited in the Gene Expression Omnibus (GEO) under accession codes GSE299180 and GSE299181.

