## Supplementary figures and tables for "CTCF is crucial for decidualization of uterine stromal cells through facilitating HOXA11 expression"

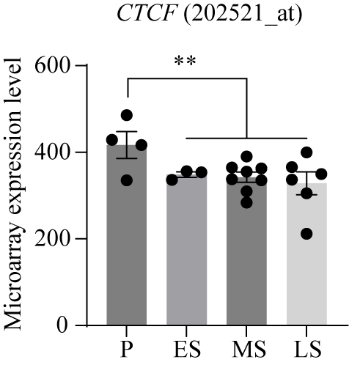

**Supplementary Figure 1. CTCF mRNA expression in the human endometrium across the menstrual cycle, as depicted in microarray dataset GSE4888.** P, proliferative phase; ES, early secretory phase; MS, middle secretory phase; LS, late secretory phase. Data are presented as means ± SEM. **, P < 0.01.

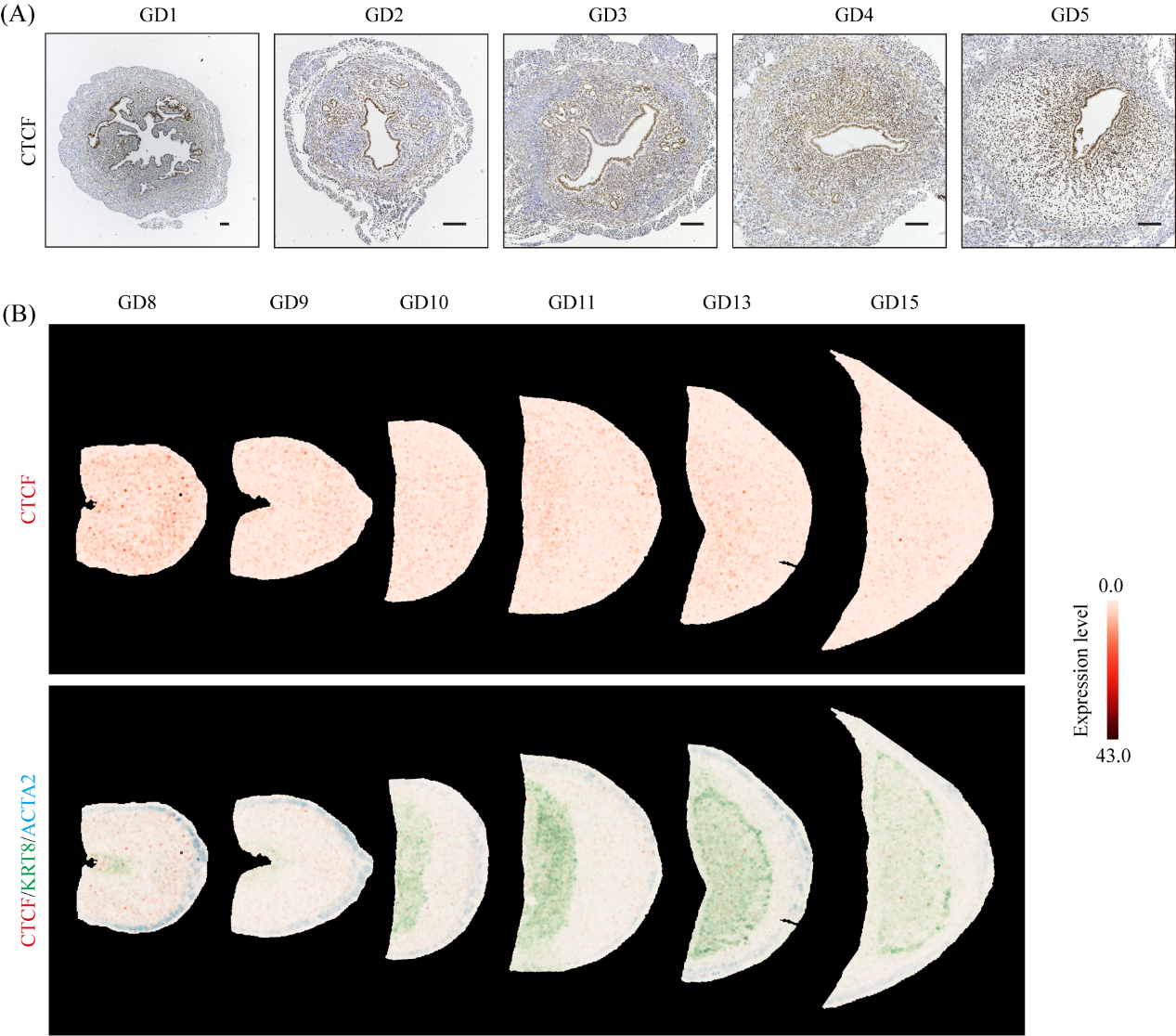

**Supplementary Figure 2. CTCF expression dynamics in the mouse uterus during pregnancy.** (A) Immunohistochemical staining reveals CTCF protein localization in the mouse uterus during the peri-implantation phase. Brown color denotes positive CTCF staining. GD, gestational day, with day 1 marking plug detection. Bar=100 μm. (B) Stereo-seq dataset analysis depicts CTCF expression trend in the mesometrial decidua or decidua basalis from GD8 to GD15 (https://db.cngb.org/stomics/mpsta/spatial/).

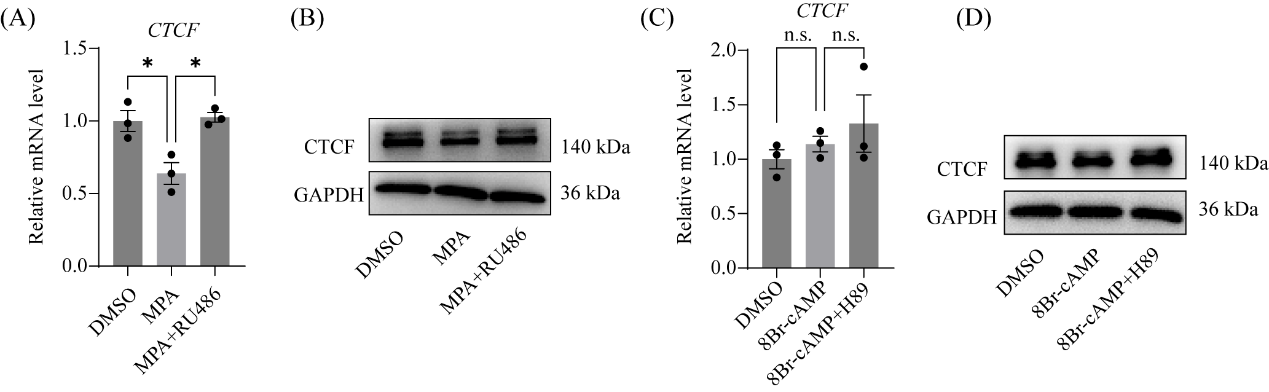

**Supplementary Figure 3. Regulatory effects of MPA and 8Br-cAMP on CTCF expression.** (A-B) CTCF mRNA (A) and protein (B) levels in hESCs following 2-day treatment with MPA alone or in combination with RU486. Data are presented as mean ± SEM. *, P < 0.05. (C-D) CTCF mRNA (C) and protein (D) expression in hESCs after 2-day treatment with 8Br-cAMP alone or co-treated with H89. Data are presented as mean ± SEM. n.s., not significant.

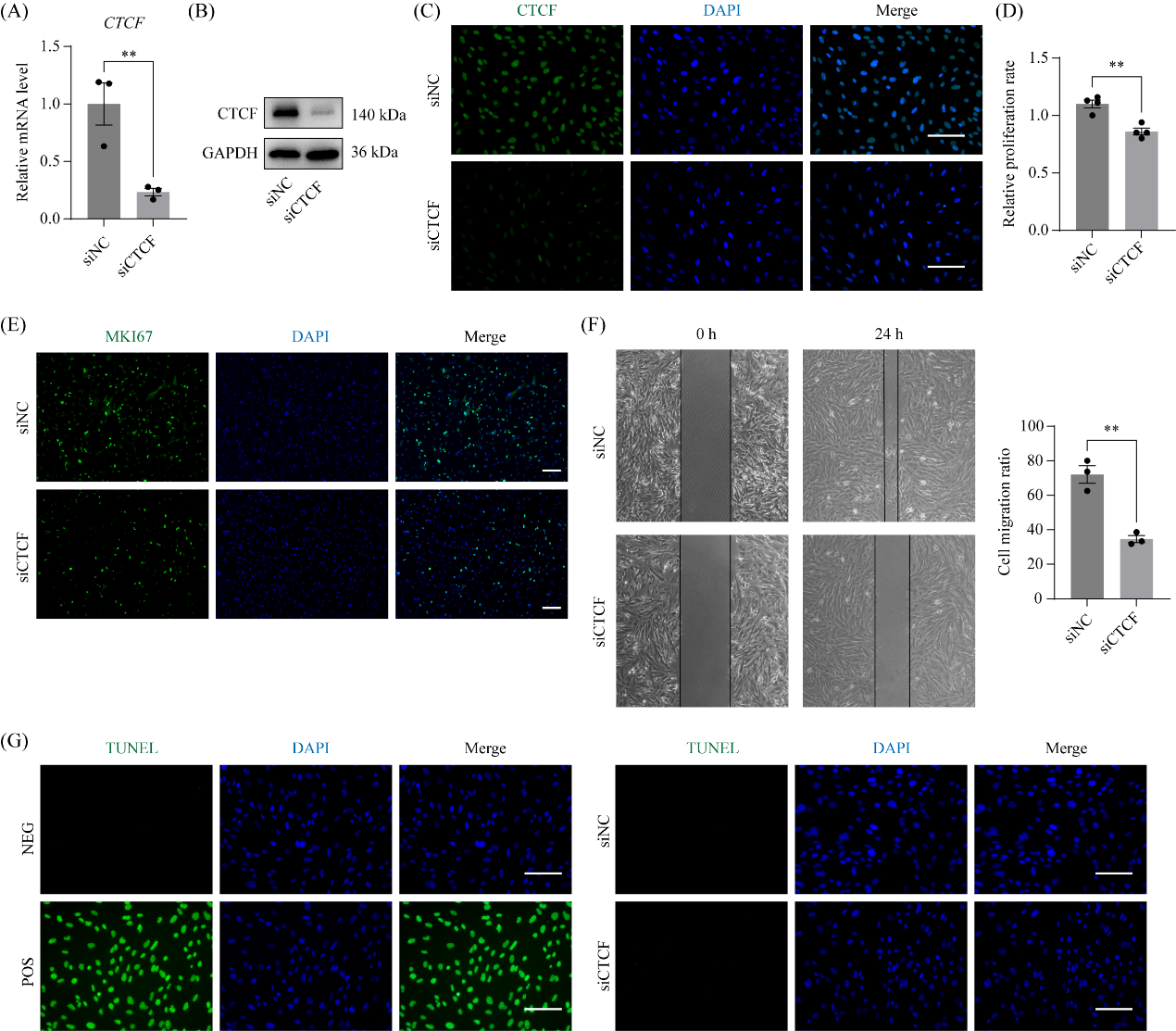

**Supplementary Figure 4. CTCF knockdown in hESCs attenuates cell proliferation and migration, with no impact on apoptosis.** (A-C) Validation of CTCF-targeting siRNA in hESCs. (A) RNA levels of CTCF post-48h siRNA-mediated knockdown. Data are presented as mean ± SEM. **, P < 0.01. (B) Western blot assessment of CTCF after 48h siRNA-mediated knockdown. (C) Immunofluorescent staining of CTCF protein in hESCs following 48h siRNA-mediated knockdown. Bar=100 μm. (D) Bar plot illustrates alterations in cell proliferation rate post-48h siRNA-mediated CTCF knockdown, measured using the CCK8 kit. Data are presented as mean ± SEM. **, P < 0.01. (E) Immunofluorescent staining of MKI67 protein in hESCs after 48h siRNA-mediated CTCF knockdown. Bar=100 μm. (F) Scratch migration assay of hESCs subjected to 24h siRNA-mediated CTCF knockdown. Data are presented as mean ± SEM. **, P < 0.01. (G) TUNEL staining evaluates cell apoptosis post-48h siRNA-mediated CTCF knockdown. NEG, no enzyme in TUNEL reaction as negative control; POS, DNase treatment prior to TUNEL reaction as positive control.

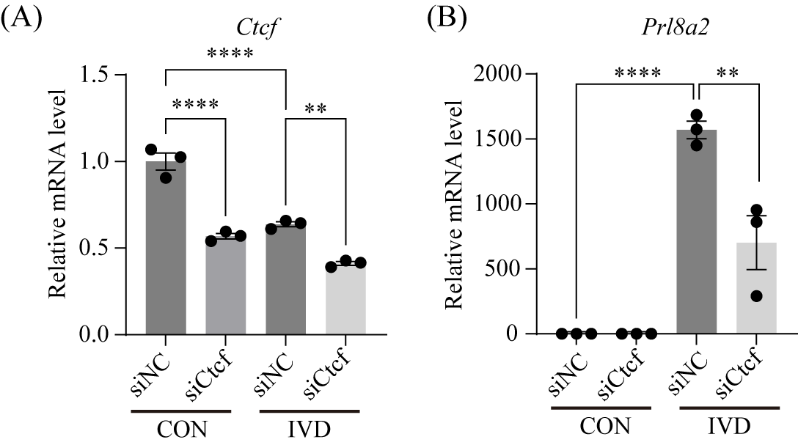

**Supplementary Figure 5. CTCF expression in mouse endometrial stromal cells (mESCs) is essential for in vitro decidualization.** Quantitative RT-PCR analysis of *Ctcf* (A) and *Prl8a2* (B) mRNA levels in primary mESCs after 3-day in vitro decidualization with CTCF knockdown. CON, vehicle control; IVD, in vitro decidualization. Data are presented as means ± SEM. **, P < 0.01; ****, P < 0.0001.

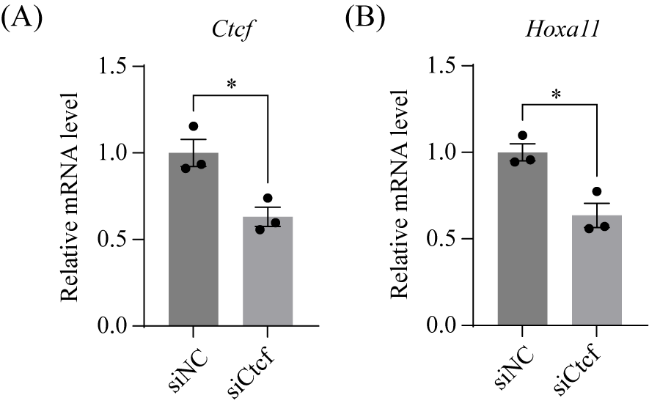

**Supplementary Figure 6. HOXA11 is positively regulated by CTCF in primary mouse endometrial stromal cells (mESCs)**. (A) CTCF knockdown in mESCs. Data are presented as mean ± SEM. *, P < 0.05. (B) Quantification of *Hoxa11* mRNA levels in mESCs after 48h of CTCF knockdown. Data are presented as mean ± SEM. *, P < 0.05.

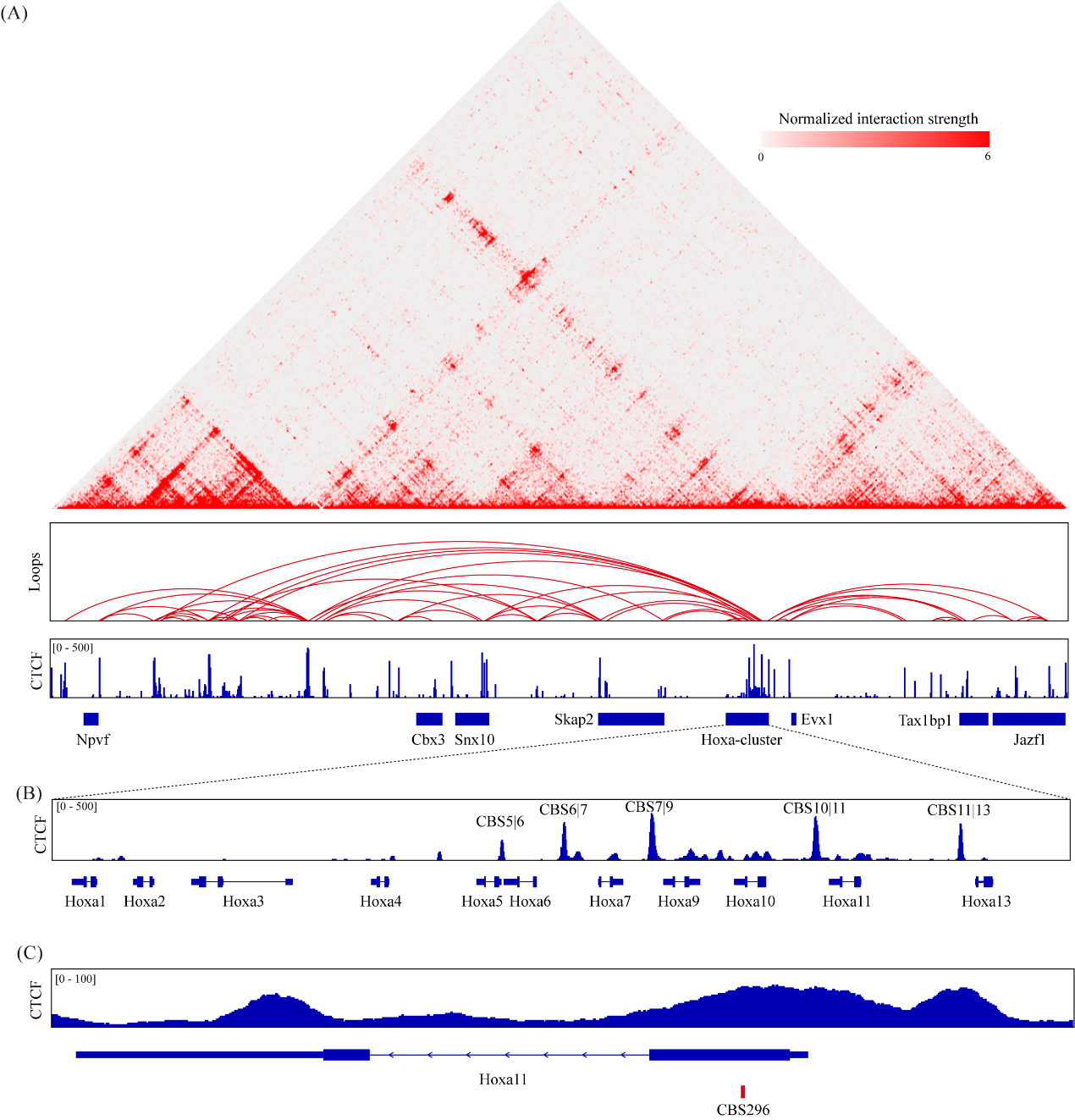

**Supplementary Figure 7. Identification of CTCF binding sites (CBSs) within the *Hoxa11* locus in the mouse uterus.** (A) Hi-C interaction map and chromatin loops depicted for the genomic region encompassing the HoxA locus in mouse uterus, derived from a public Hi-C dataset (GSM3587145). (B) IGV (Integrative Genomics Viewer) screenshots display CTCF ChIP-seq signal enrichment at the HoxA locus in mouse uterus, obtained from a public CTCF ChIP-seq dataset (GSM7102075). (C) Discovery of a novel CTCF binding site (CBS296) proximal to the TSS of *Hoxa11* using PWMScan (https://epd.expasy.org/pwmtools/cgi-bin/pwmtools/pwmscan_form_parser.cgi).

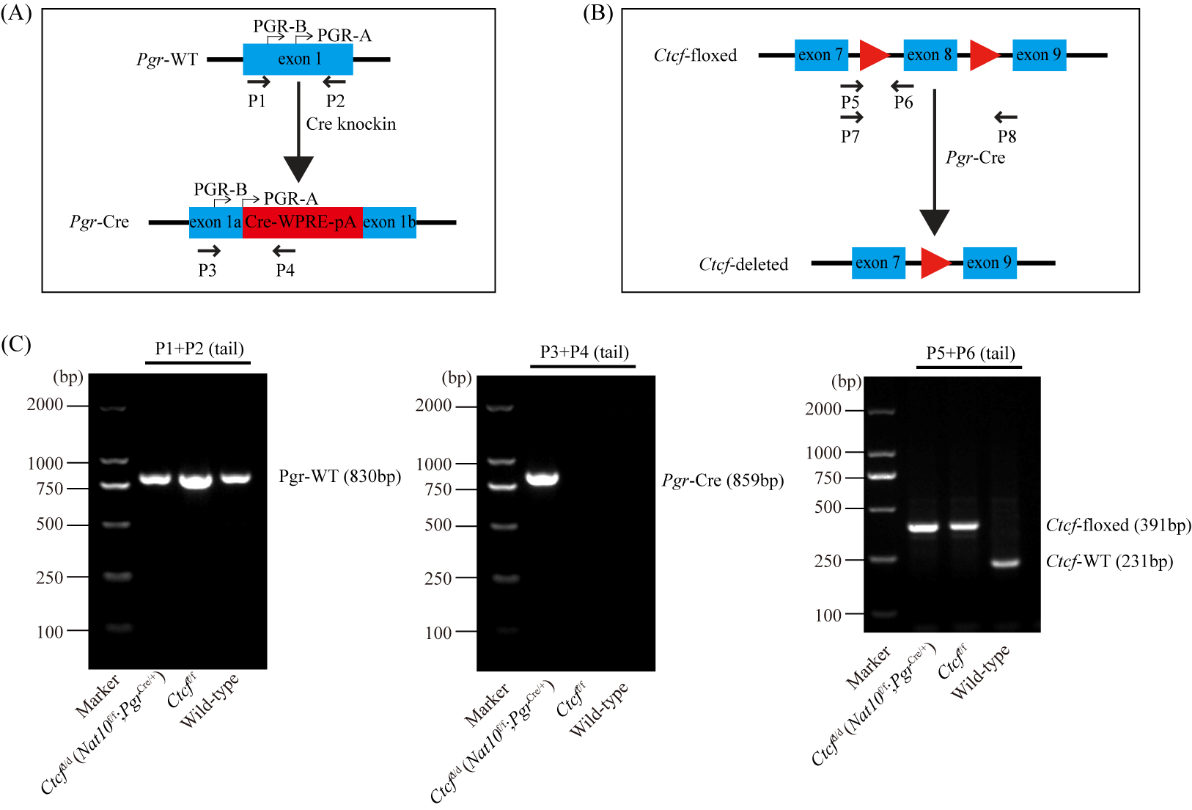

**Supplementary Figure 8. Genotyping assessment of *Ctcf*^d/d^ and *Ctcf*^f/f^ mice.** (A) Schematic representation of genotyping primers targeting the *Pgr*-Cre allele. (B) Illustration of genotyping primers for the *Ctcf*-floxed allele. P7/P8 was utilized in Fig. 5C. (C) Genotyping PCR results for *Ctcf*^d/d^ and *Ctcf*^f/f^ mice.

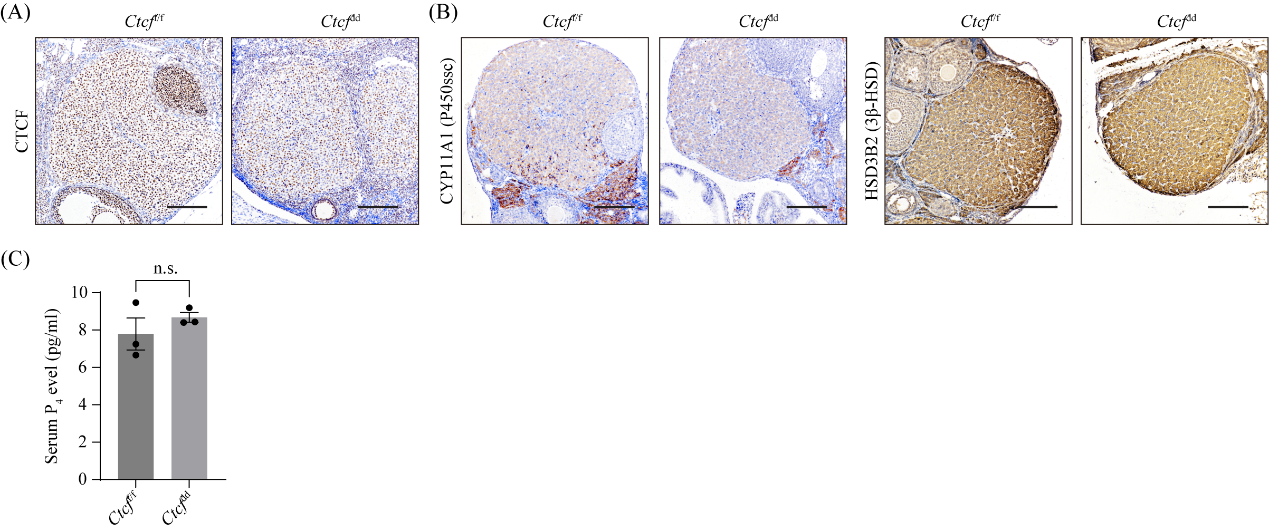

**Supplementary Figure 9. Ovarian function in *Ctcf*^d/d^ mice remains normal.** (A) Immunohistochemical staining for CTCF in the ovaries of *Ctcf*^d/d^ and *Ctcf*^f/f^ mice on GD4. Bar=100 μm. (B) Immunohistochemical visualization of CYP11A1 and HSD3B2 in the ovaries of *Ctcf*^d/d^ and *Ctcf*^f/f^ mice on GD4. Bar=100 μm. (C) Circulating P_4_ levels in *Ctcf*^d/d^ and *Ctcf*^f/f^ mice on GD4, measured using ELISA kits (Genkern, Guangzhou, China) as per the manufacturer's guidelines. n.s., not significant.

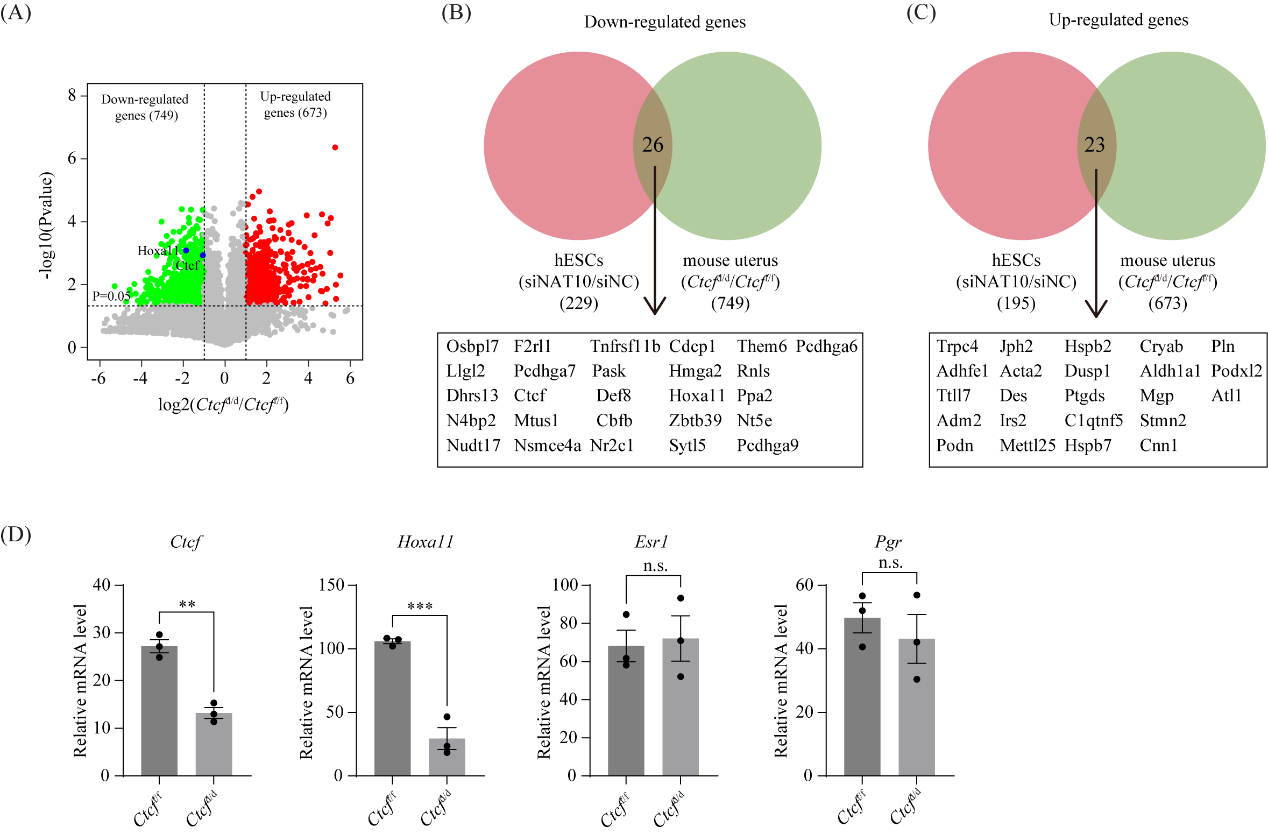

**Supplementary Figure 10. RNA-seq analysis reveals that *Hoxa11* mRNA was down-regulated in *Ctcf*^d/d^ uterus on GD4.** (A) Volcano plot for differentially expressed genes in the uterus from *Ctcf*^d/d^ mice and *Ctcf*^f/f^ mice on GD4 from RNA-seq analysis. (B-C) Venn diagram depicting the overlap of differentially expressed genes identified in CTCF-knockdown hESCs and *Ctcf*-knockout mouse uteri. (D) Bar plot showing the expression of *Ctcf*, *Hoxa11*, *Esr1* and *Pgr* in the uterus of *Ctcf*^d/d^ mice compared to *Ctcf*^f/f^ mice, based on RNA-seq data. **, P < 0.01; ***, P < 0.001.

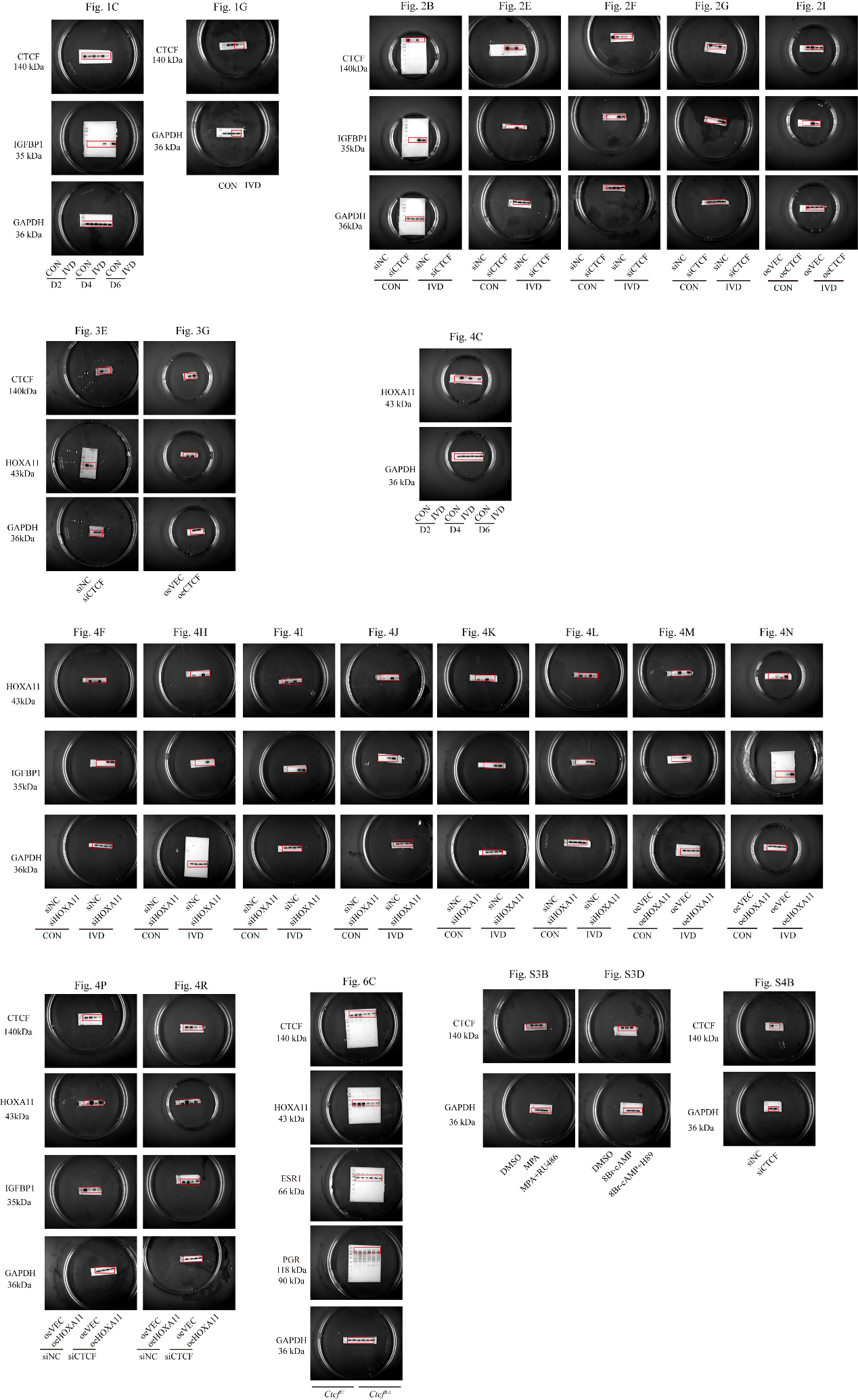

**Supplementary Figure 11. Uncropped images of the western blots presented.**

**Supplementary Tables**

**Supplementary Table 1. Differentially expressed genes in CTCF-knockdown human endometrial stromal cells (hESCs) versus controls (fold change > 2 and p-value < 0.05).**

**Supplementary Table 2. Differentially expressed genes in CTCF knockout uteri compared to controls on gestation day 4 (GD4) (fold change > 2 and p-value < 0.05).**

**Supplementary Table 3. Detailed information of human participants for isolation of primary human endometrial stromal cells (hESCs) in this study.**

**Supplementary Table 4. Primers used in this study.**

**Supplementary Table 5. Antibodies used in this study.**

**Supplementary Table 6. siRNAs and overexpression vectors used in this study.**
